# Mapping Structural Drivers of Insulin and its Analogs at the IGF-1 Receptor Using Molecular Dynamics and Free Energy Calculations

**DOI:** 10.1101/2023.12.02.569705

**Authors:** Mohan Maruthi Sena, Ramakrishnan C, M. Michael Gromiha, Monalisa Chatterji, Anand Khedkar, Anirudh Ranganathan

## Abstract

Insulin and insulin-like growth factor-1 receptors (IR, IGF-1R) belong to the family of receptor tyrosine kinases (RTKs), and share close structural resemblance. However, these receptors exhibit distinct activity profiles and functions in vivo. Binding of insulin to IGF-1R results in additional growth-factor-like behavior and cell proliferation, but its ∼100-fold reduced affinity to IGF-1R limits off-target activity. However, insulin analogs with increased potency at IGF-1R have oncogenicity as a key safety concern. Hence, the ability to accurately predict potency of novel analogs at IGF-1R could represent a key breakthrough towards rational insulin design. To date, a comprehensive molecular level understanding of insulin interactions at IGF-1R has remained elusive. This study capitalized on recent advancements in structural biology that provided high resolution structures of IGF-1R bound to IGF-1 and insulin. Initially, molecular dynamics (MD) simulations were employed to unravel the intricate interactions that characterize the receptor-ligand pairs. Next, free energy perturbation (FEP) calculations were performed to understand the increased affinity observed in insulin analogs, X10 and glargine. Subsequently, multiple mutations at the B10 position of insulin spanning different activities at IGF-1R and different metabolites of insulin glargine, encompassing various mitogenic potencies were studied using FEP. The calculations successfully captured directional shifts in potency for all studied mutants, with approximately 50% of the predicted values falling within 1 kcal/mol of experiment. Beyond its impressive accuracy, FEP’s ability to provide a detailed understanding of protein- and solvent-mediated contributions to the observed functional profiles underscores its utility in designing safe IGF-1R selective novel insulin analogs.

## Introduction

The type 1 insulin-like growth factor receptor (IGF-1R) is responsible for various cellular processes such as proliferation, differentiation, survival, and metabolism.^1–4^ These biological functions are mediated by the binding of cognate peptide hormone IGF-1.^5,6^ This peptide is synthesized primarily in the liver in response to endocrine growth hormone (GH) stimulus. However, multiple tissues secrete this hormone for autocrine and paracrine purposes.^7^

IGF-1 is a polypeptide with 70 amino acids, which binds to IGF-1R to primarily activate the Akt pathway.^8,9^ IGF-1 shares significant similarity to insulin (∼50%) and IGF-II (60%).^6^ The cognate receptor IGF-1R belongs to receptor tyrosine kinase (RTK) family of proteins that also includes IR (insulin receptor) and growth factor IGF-2R, orphan insulin related receptor.^2,10^ RTK’s receptors exhibit a shared structural framework comprising of an extracellular domain (ECD), a transmembrane segment and an intracellular kinase domain.^11^ In line with their endogenous ligands, Insulin receptor (IR) and IGF-1R shares 50 % sequence homology overall and above 80 % sequence homology in the intracellular region.^12^ Despite this high degree of similarity between the receptors and commonalities in intracellular signaling, they perform distinct biological functions.^7,13^ Whereas IR activation is primarily responsible for glucose metabolism, IGF-1R signaling is associated with cell proliferation.^2,8,10,13–15^ However, due to the high levels of similarity between both receptor and ligand sequences, insulin can bind and fully activate the IGF-1R, albeit with a 100-fold reduced potency when compared to the cognate IGF-1.^16^ The ability of endogenous insulin to activate IGF-1R signaling and consequently cause cell proliferation, renders this receptor a key off-target when designing and evaluating modified insulin analogs.^12,17,18^ Hence, it is desirable for insulin analogs to have a similar or greater IR/IGF-1R affinity ratio compared to human insulin to ensure that oncogenicity risks due to aberrant IGF-1R mediated cell proliferation remain low.^18,19^

Multiple insulin analogs have been well-characterized in terms of IGF-1R binding, and mitogenic potency.^20–23^ However a structural understanding of the drivers of ligand engagement at this receptor remains elusive. Efforts towards elucidating a molecular-level understanding of ligand recognition at IGF-1R has been hamstrung by the lack of high-resolution structures. Recently, a breakthrough has been achieved in terms of ligand-bound IGF-1R structures. Xu *et.al*, provided the first high-resolution of the IGF-1R ectodomain structure showing the apo structure exists in “Λ” (inverted V-shape) state.^2^ Upon IGF-1 binding the receptor shifts conformation towards an activated state which has separation observed between L1-CR domain away from Fn-III-2’.^2^ The binding site lies between with the Fn-III-2’, L1 and α-CT of the receptor in the activated state. During the course of this work, multiple cryo-EM or X-ray diffraction derived structures were obtained for IGF-1R in complex with IGF-1 and insulin (Table S1). Interestingly a new cryo-EM structure of insulin/IGF-1 bound IGF-1R structure was elucidated by Zhang *et al.* without any deletion, mutations or truncations of the receptor ectodomain (ECD).^10^ In line, with the first structure of the IGF-1:IGF-1R complex, only one insulin molecule was observed to bind to the receptor, which is in contrast to the insulin:IR complexes, where up to 4 insulin molecules can bind to the IR-homodimer.^24^ However, there were also notable differences between the cryo-EM structure and previously reported structures. One such difference was observed in the distance between two Fn-III-3 domains which decreased from 120 Å to 80 Å and the interactions of L1 domain with Fn-III-1 domain rather than Fn-III-2 which were reported previously.^2^ The cryo-EM structure displayed an active state conformation of IGF-1R, which was more in-line with that observed for the IR.^10^ In this study, we have used structures from both Xu *et al.* (X-ray crystallography) and Zhang *et al.* (Cryo-EM) to understand ligand engagement at IGF-1R.^2,10^ If these structures, could be leveraged to provide a clear understanding of the key molecular determinants of ligand engagement at IGF-1R it could have a significant impact on designing safer insulin analogs with large factors-of-safety.

An interesting analog of insulin which attracted lot of attention due to its fast acting property (due to reduction of self-association) and increased metabolic potency was insulin X10 (also referred to as B10Asp).^19,21,22,25^ A major drawback of this analog was from the perspective of safety since it increased mitogenic potency 3-20 fold compared to human insulin (primarily via increased IGF-1R binding).^22,26^ Insulin X10 carries a single change from the human insulin sequence, where the naturally occurring histidine at position B10 is replaced with an aspartic acid.^20,21,27^ The high mitogenic potency exhibited by insulin X10, unfortunately translated to cancers upon prolonged exposure in rats.^19,28^ This led to its withdrawal from phase-2 clinical trials.^19,28,29^ Different amino acid substitutions at B10 result in a spread of mitogenic potencies.^22^ Hence, insulin X10 and other B10 variants of human insulin presented a highly interesting molecule set for this study.

Exogenous administration of modified insulins (analogs) or formulated human insulin with different pharmacokinetic/pharmacodynamic profiles are used to provide sufficient glycemic control in the treatment of diabetes.^20–22,30,31^ These insulin analogs in current use for the management of type I or II diabetes have been well studied in terms of their metabolic and mitogenic potencies mediated primarily via their action on the target IR, and off-target IGF-1R.^20–23,31^ One such analog, insulin glargine (LANTUS®), is a long-acting insulin, which offers glycemic control for ∼24 hours.^23,32^ Glargine has been available for diabetes treatment for more than two decades and carries the following modifications when compared to human insulin^23,33,34^ - (i) The replacement of the C-terminal Asparagine of the A-chain with a glycine (A21Gly) (ii) Extension of the insulin B-chain of human insulin with two arginines (B31Arg/B32Arg). Interestingly, glargine has a significantly higher mitogenic potency (and IGF-1R affinity), when compared to human insulin, while additionally possessing a lower metabolic potency (and IR affinity).^23^ Hence, its target/off-target (IR/IGF-1R) potency ratio is significantly lower than human insulin, resulting in a potentially lowered safety window. However, insulin glargine has a complex dissociation profile following sub-cutaneous administration due to its increased pI of close to the physiological pH of 7 and is processed into a series of intermediates and metabolites due to enzymatic cleavage.^23,35^ This combination of factors allows for sufficient safety of the molecule.^23^ The significantly enhanced IGF-1R affinity of insulin glargine and the presence of a series of intermediates/metabolites with varied mitogenic profiles rendered this series of molecules an ideal test set for the current study.

Since the structure of insulin itself was elucidated decades ago, a majority of computational studies were carried out explaining the conformational dynamics of monomeric, dimeric and hexmeric form of insulin.^36–39^ A similar lack of high-resolution structures for insulin:IR hamstrung computational efforts, limiting the field to a few studies that relied on ligand docking to the apo IR structure, followed by MD simulations.^40–42^ The structures have since revealed significant differences between the computationally predicted bound conformation and the actual insulin binding mode at IR.^40,41,43^ Whereas, there were multiple computational studies on free insulin in an attempt to elucidate the bioactive conformation, efforts on IGF-1 was restricted to the work of Papaioannou *et.al* who similarly predicted that IGF-1 binding required the opening of BC-CT (B chain-C-terminal) similar to insulin, and additional opening of BC-CT away from the C-domain.^44^ In fact, our work, to the best of our knowledge represents the first structure-based study involving IGF-1R. Hence, we defined three primary goals to this study. The first was to provide a framework to understand the stability and interaction networks characterizing insulin bound to its anti-target the IGF-1R using short MD simulations with IGF-1 bound receptor providing a base framework. Next, we wished to understand the molecular-drivers of potency jumps at IGF-1R for well-studied insulin analogs, X10 and glargine using free energy perturbation (FEP). Since, potency jumps at IGF-1R are highly undesirable from a safety perspective due to the chronic nature of diabetes treatment, this understanding could provide key insights towards the avoidance of off-target binding, with insulin analogs. Finally, we attempted to provide an enhanced understanding of the different potency profiles exhibited by closely related analogs/metabolites of insulin X10 and insulin glargine. If the methods used here could provide an accurate readout for IGF-1R potency that would represent a key breakthrough towards the use of FEP as a generalized design tool for safer novel insulins. The deconvolution of binding energetics and an enhanced understanding of the structural drivers of recognition at the off-target could have implications for design of selective peptide ligands beyond insulins.

## RESULTS

### Analysis of the IGF-1R structure used in this study

IGF-1R is an functional assembly of two α-chains and two β-chains forming a heterodimer (αβ)_2_.^2,45^ ^10^ The α-chain constitutes of L1 (Leucine rich), L2 (Leucine rich), CR (Cysteine-rich) domains besides two fibronectin-like segments Fn-III-1, and few residues of Fn-III-2 whereas the β-chain contains, the remainder of Fn-III-2, Fn-III-3, a single-spanning transmembrane (TM) segment, and an intracellular tyrosine kinase (TK) domain.^2,10^ The L1, CR, FnIII-1-3 segments together constitute the extracellular domain (ECD) of the IGF-1R and contains the ligand binding site(s).^5^ Recently, EM and XRD have provided high resolution structures of cognate and non-cognate ligands bound to the to the ECD including recent studies that provided the complete ECD bound to Insulin/IGF-1 and without any mutations or deletions in the flexible regions of the receptor.^2,10^ Whereas, both ligand bound structures (Insulin/IGF-1) had overall similarities, there were also differences observed. The major distinction involved conformational changes in the receptor upon ligand binding, which showed the expected transition from an inverted “V” shape to a T-shape in the EM (insulin-bound) structure [PDB ID: 6JK8] but not in the XRD (IGF-1 bound) structure [PDB ID: 5U8Q]. ^2,10^ A recent unpublished electron microscopy (EM) structure by Xi *et al.,* with the structure deposited in the [PDB ID: 7YRR], showed the IGF-1R in complex with two IGF-1 molecules. Here, the receptor adopts an inverted “V” shape, similar to the XRD structure of IGF-1 bound to IGF1-R by Xu *et al* (PDB ID: 5U8Q). On the other hand, Zhang *et al.* observed a transition to a “T”-like conformation upon IGF-1 binding, albeit with a lower resolution EM structure. Hence, given the existence of IGF-1R in two distinct ligand-bound conformations, we used ligand-bound structures from Xu *et al.* (XRD) and Zhang *et al.* (EM) as reference structures in this study.^2,10^

### Comparison of XRD and EM structure

The following differences were observed between the XRD structure of Xu *et al*. and EM structure of Zhang *et al.*^2,10^ In both studies, the apo-state adopts an inverted “V” shape, while the electron microscopy (EM) structure reveals that the conformation transitions to an active T-shape upon insulin binding.^2,10^ In the XRD structure presented by Xu *et al.,* obtaining the crystal structure of the complex involved the removal of several flexible domains, which led to a partial loss of functional architecture.^10^ Additionally, only one insulin is bound to receptor in the EM study but XRD study observed two IGF-1 molecules bound at saturating concentrations.^2,10^ In both the studies, L1-CR-L2 domains constitute the hormone binding regions and Fn-III-2/3 constituted the legs spanning to the transmembrane region. Since, as the IGF-1R exists as dimer (αβ)_2_, the monomer (αβ) which is hormone free has significant conformational changes compared to hormone bound region.^10^ It was also observed that the α-CT’ present in hormone free monomer (αβ) interacts with L’ which was not observed in the earlier reported structures from Xu *et al*.^2,10^ The overall conformation of the insulin/IGF-1 in the XRD/EM structures remain same but in the EM structure, insulin mediates interactions between L1-CR-L2 (binding region), Fn-III-1’ and α-CT, whereas IGF-1 interacts with L1-CR, α-CT’ and Fn-III-1’.^10^ The variation in placement of the α-CT domain upon ligand binding can be observed via its interactions with 23Phe, 24Tyr, 25Phe, 26Asn of residues of IGF-1 with 701Phe, 702Val, 704Arg of α-CT in the XRD structure and equivalent 735Arg, 560Lys, 514Pro 721Lys with 19Tyr, 15Leu, 7Cys, 48Phe residues of insulin in the EM complex. Since, there were significant differences observed between ligand bound conformations in the two structures, we chose the EM structure of the insulin-IGF-1R complex (PDB accession code: 6JK8) and the XRD structure of the IGF-1-IGF-1R (PDB accession code: 5U8Q) complex as our reference structures.^2,10^ We used these reference structures to assess insulin glargine and its metabolites, and various substitutions at X10.

### Short MD simulations of the IGF-1R with WT-ins/IGF-1 to assess stability and ligand engagement

To understand and analyze the interactions of the cognate and non-cognate ligands with IGF-1R, all-atom molecular dynamics (MD) simulations were performed for fully solvated system of insulin/IGF-1-IGF-1R complex at 300 K. The starting coordinates for the complex were obtained from Xu *et.al*., [PDB accession code-5U8Q] and Zhang *et.al.,* [PDB accession code- 6JK8].^2,46^ Since bound insulin to IGF-1R receptor was not available initially in the conformation represented by XRD (IGF-1 bound) structure, homology modeling were used to get insulin bound IGF-1R structure in this conformation. This was done to understand the behavior of insulin-IGF-1R complexes in both observed ligand-bound conformations of this receptor. MD simulations were performed by initially applying restraints on the heavy atoms of the protein-ligand system and later simulated with complete flexibility.

The RMSD obtained for the cognate/non-cognate complexes is shown in Figure 2a, which indicates the overall stability of the respective complexes throughout the simulation time considered. The RMSD of cognate complex and non-cognate complex using XRD structure are comparatively lower in comparison to the RMSD of the non-cognate complex using EM structure. The higher RMSD of EM structure can be attributed to following structural insights, the higher flexible regions of helical turn of A-chain residues (A1-A8) and B-chain residues (B14-B20) compared to initial structure (Figure 2b). This observed loss of structure is due to the interactions of these residues with the residues of L1-domain. Additionally solvent accessible surface area (SASA) were calculated for each ligand and thus obtained values are in line with experimentally reported surface area values (Table S2).^24^

To further understand overall stability of bound complexes, intermolecular interactions between the ligand and receptor were analyzed. In order to provide a simplistic framework for understanding the differences in intermolecular interactions were divided into WT-ins and IGF-1 (ligands) into sub-segments based on secondary structure elements (Figure 2d). When an occupancy cut-off of 90% was applied to interactions within a 4.5 Å shell around the ligand, the analysis revealed key interactions with receptor which contributed towards overall stability of the complex (Figure 2d). In all three complexes, highly stable interactions are dominated by the C-terminal of A-chain of the respective ligands with the α-CT domain of the receptor. Additionally, the C-terminal of the B-chain of the IGF-1 and insulin bound in the global XRD structure conformation possess stable interactions with receptor which was not observed in the EM structure. In all the three cases, the A19Tyr residue of insulin and equivalent (A21Tyr) residue in IGF-1 interacts with the α-CT domain of the receptor. Binding of non-cognate ligand insulin to IGF-1R resulted in the distinct patterns of interaction. In XRD and EM, there were some common notable interactions observed between insulin and receptor, eg. B17Leu interacts with 1573Phe of Fn-III-1 domain. At the same time, there were distinct interactions observed at the N-terminal of the insulin B-chain, when starting from the EM structure which was not observed when initating from the XRD structure and vice-versa eg. A3Val interact with 694Asn whereas in EM it interacts with 695Phe. Overall, the variation in interaction patterns between the insulin-IGF1R complexes derived from XRD/EM structures can largely be attributed to changes in the overall conformation of domains around the ligand. These interaction patterns in both the structures reveal potential key interactions for the observed ligand bound conformations of the IGF-1R and enhance our understanding of peptide binding to this receptor.

### Understanding the increase in potency of super active X10 insulin

The B10 position plays a crucial role in the hexamerisation of human insulin.^47–49^ Substitution of Histidine at B10 position in WT-ins by Aspartic acid has attracted lot of attention owing to its spike in metabolic potency at IR compared to human insulin, for which it was labeled as “superactive insulin”.^22,27,50,51^ It has also been observed that this analog has enhanced mitogenic potency, which exceeds its increase in metabolic potency.^20,22,23,52–54^ Analysis of B10 with static experimental structures have shed limited light on the molecular origin of these effects due to the absence of any direct interactions for B10 with IGF-1R.^2^ FEP was employed to investigate the impact of multiple experimentally-assessed substitutions at the B10 position, aiming to enhance comprehension of the association between the nature of amino acid at this position and its overall impact on the functionality. FEP calculations were performed using the final snapshot obtained from the equilibration step of MD simulations performed in the previous step. All the FEP calculations were carried out using spherical boundary conditions in which the ligand-bound state is closer to the starting structure (Figure 1b, Figure S1a). The simulation process involved allowing all atoms located within a specified sphere to move freely while tightly restraining the residues located outside the sphere to their initial coordinates. This approach helped to maintain the overall structure of the receptor intact while studying the region of interest within the sphere. The optimal sphere radii were chosen to ensure such that all the necessary domains for function are included. Throughout the simulations, no significant fluctuations in the structure within the sphere were observed, and the RMSD values remained < 0.3 Å. Since, free energy calculations are more sensitive to the initial conformation of the protein of interest both EM [PDB ID: 6JK8] and XRD [PDB ID: 5U8Q] derived structures were considered for the HisB10Asp mutation.^2,10^ In total 1.3 μs simulations were performed across different B10 variants with ∼ 220 ns used for each FEP calculation at the receptor. To determine the relative binding free energies (ΔΔG_bind_) of each mutant, thermodynamic cycle was constructed, with free ligand in water serving as the reference state (Figure S1b). The resulting ΔΔG_bind_ values were then compared to that of the WT-ins to measure the relative affinities of each molecule at the IGF-1R.

**Figure 1:**
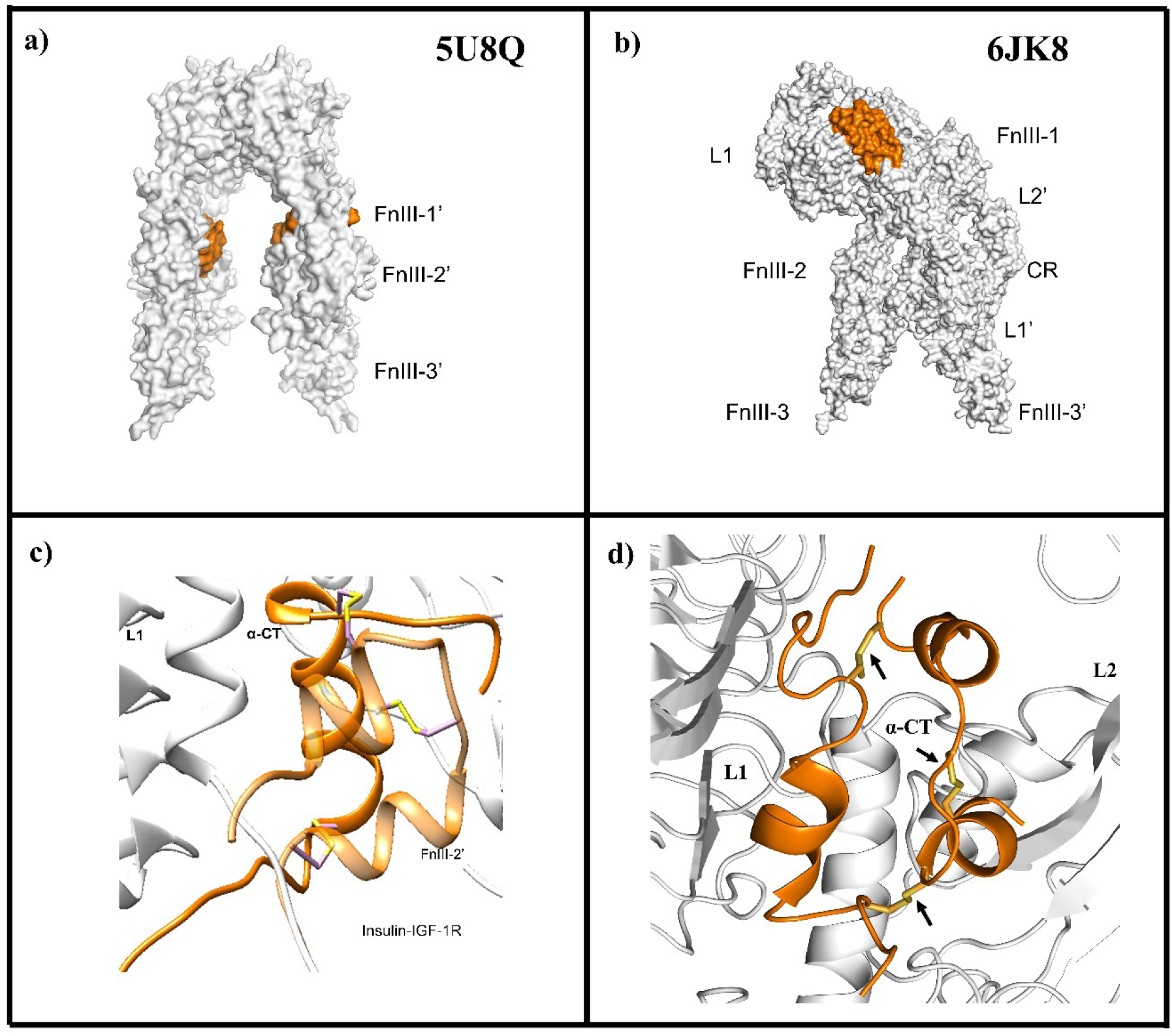
Structures of experimentally obtained IGF-1R structures a) XRD (5U8Q) and b) EM (6JK8) used in this study. In the top panel, the surface representations of the IGF-1R receptors are shown with their respective ’V’ conformations (from XRD) and the active ’T’ shape conformation (from EM). The binding sites for the receptor are highlighted in orange. In the bottom panel (c,d), a zoomed-in view of XRD, EM structure of the insulin binding with the receptor is presented respectively, with the insulin molecule shown as an orange cartoon. The arrows indicate the intact inter- and intra-chain disulfide bonds in the complex.

**Figure 2:**
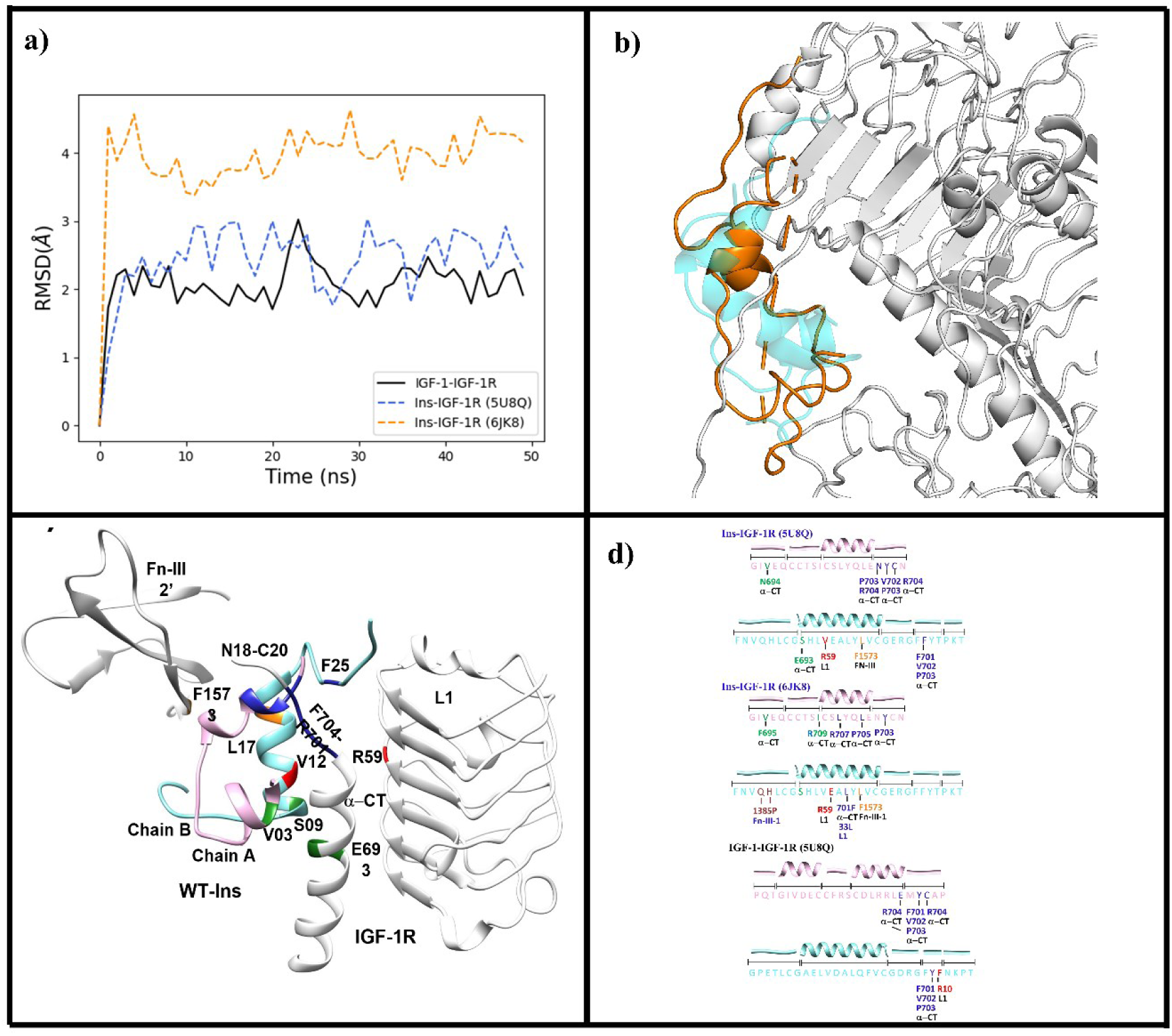
Assessment of stability of insulin-non-cognate, IGF-1-cognate complexes from MD simulations with the starting coordinates as reference a) Cα-RMSD of the three simulation conditions with respect to the initial structure, IGF-1-IGF-1R (black solid line), insulin-IGF-1R (blue, dashed line, 5U8Q), and insulin-IGF-1R (orange, dashed line, 6JK8), b) insulin-IGF-1R complex: the insulin average structure from XRD (cyan cartoon) is shown in alignment with the average structure of insulin (orange cartoon) and receptor (grey cartoon) obtained from MD simulations. c) Analysis of stable contacts of WT-ins in IGF-1R receptors with more than 90% lifetime obtained from MD simulations. Ligand-receptor contacts were defined using a distance cutoff of 4.5 Å from any ligand residue. The respective receptors in IGF-1R are represented as grey cartoons, the chain-A or chain-A equivalent of the respective ligands WT-ins or IGF-1: cyan and the chain-B or chain-B equivalent as Mauve. The interacting residues from ligand and receptor are identically color-coded domain-wise and domain names/residue names are further highlighted in text, d) shows the sequence view of WT-ins and the resolved portion of IGF-1. The interactions of ligand residues with specific receptor residues in the cognate and non-cognate complexes, namely, WT-ins-IR (5U8Q) (top), WT-ins-IGF-1R (6JK8) (middle), IGF-1:IGF-1R (bottom), are shown in text. For convenience IGF-1 is shown as two chains to allow for easy comparison. The interacting residues are color-coded domain-wise using the same scheme as used in the structures (c).

The ΔΔG_bind_ calculated for B10Asp with XRD structure is -2.9 kcal/mol and with EM structure it is -3.46 kcal/mol, showing good consistency irrespective of the starting conformation. This allowed the HisB10Asp mutation to serve as a bridging calculation, for performing different mutations using either the XRD or EM structures. The experimental ΔΔG_bind_ calculated from various experimental studies were -1.26 kcal/mol, showing that FEP captured the gain-of-function associated with this mutation, albeit with an overestimation of magnitude. The decomposition of ΔΔG_bind_ revealed that the large increase in affinity that has been observed for the B10Asp variant was driven by electrostatics. Structural analysis revealed no direct interactions with the receptor in either WT-ins or B10Asp. To understand the molecular origins of such a large gain in potency at the IGF-1R we turned to solvent interaction network analysis from large-scale MD simulations of the end states (B10His & B10Asp). The analysis highlighted a notable distinction in the receptor-ligand interactions mediated by water molecules between WT-ins and B10Asp variants at IGF-1R (Figure 4, Table S3). Substituting histidine with aspartic acid at B10 led to significant modifications in the first- and second-order water-mediated contacts between the receptor and ligand (Table S3). The occupancy of aspartic acid at B10 resulted in an approximately 300% rise in water-mediated contacts to the receptor, when compared to the native histidine (Figure 4 and Table S3). Additionally, these interactions extended over different domains of the receptors, serving as a bridge that stabilized the entire ligand-bound conformations of the IGF-1R.

**Figure 4:**
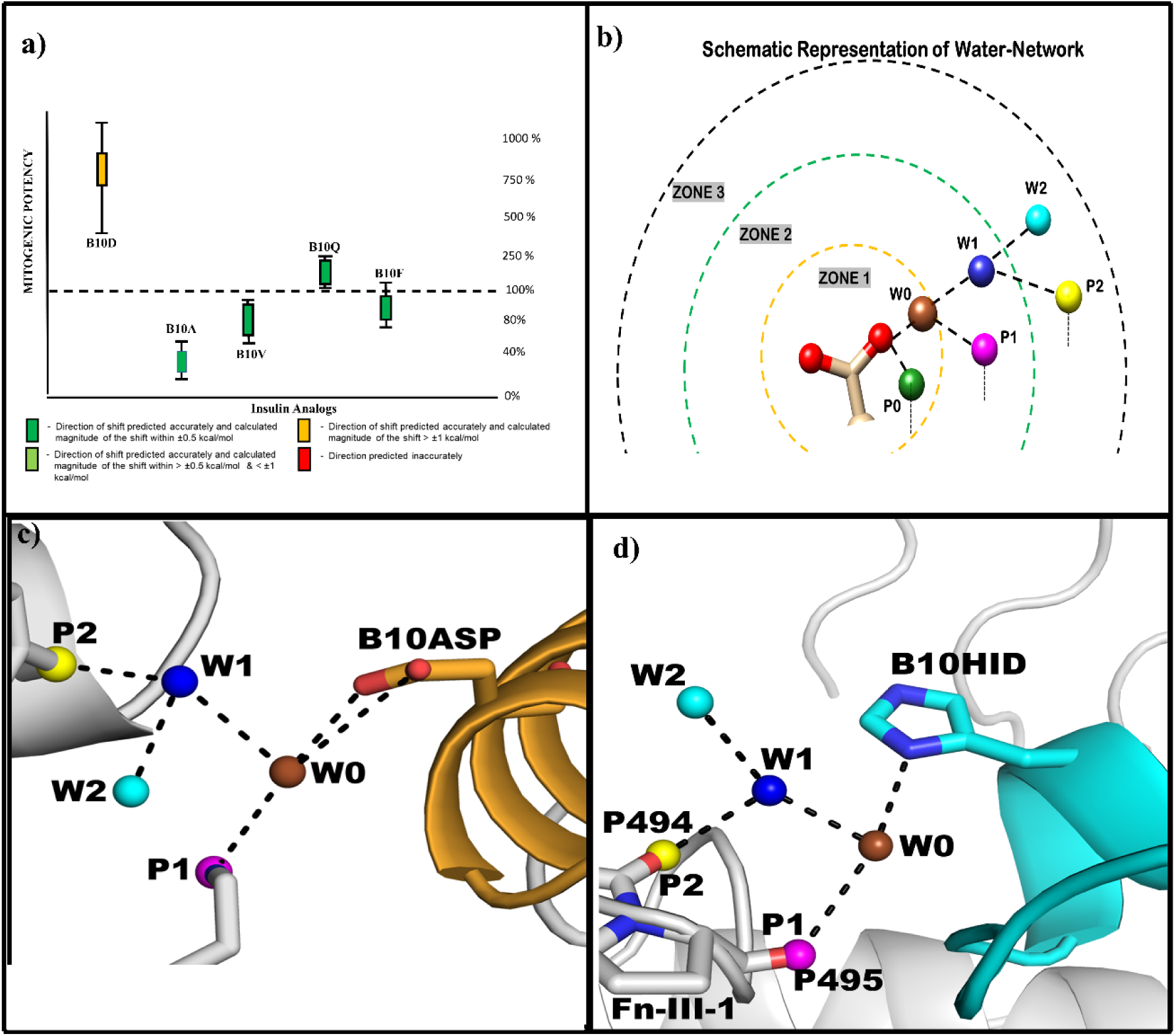
Analysis of the water-mediated interaction network around the position B10 of insulin in IGF-1R. a) Correlation plot between mitogenic potencies obtained from FEP and various experiments. b) A schematic representations of the classification of water-mediated interactions around a residue is shown. The aspartic acid residue occupying position B10 are shown in ball & stick representation, polar oxygen and nitrogen atoms are shown red or blue spheres respectively. First order or direct polar interactions with between B10 and a receptor residue is considered P0 and similarly W0 for a water contact. Second order contacts are classified as a W0-receptor polar contact defined as P1 or W0-water contact defined as W1. Similarly, third order contacts are defined as W1-receptor polar contact defined as P2 or W1-water contacts defined as W2. c) Shows the water mediated contacts for B10Asp-IGF-1R. d) Shows the water mediated contacts for B10His-IGF-1R.

Besides the evaluation of B10Asp, Glendorf *et al*. analyzed ten different single amino acid substitutions at the B10 position at IR in terms of their binding, metabolic, and mitogenic potency.^22^ In this study, we investigated four other representative amino acid mutations at B10, namely B10Ala (small), B10Val (hydrophobic alkyl), B10Phe (hydrophobic aromatic) and B10Gln (larger polar). While Glendorf *et al*. study considered DesB30-insulins, we conducted our calculations using full-length insulin to ensure consistency, given the reported high degree of correspondence between Des-B30 and full-length insulins.^55^ The calculated ΔΔG_bind_ from FEP values with full-length insulins for these analogs largely correlated with the observed experimental affinities IGF-1R (Figure 4a, Table S4).

#### B10Ala & Val

The substitution of small aliphatic hydrophobic residues such as Alanine and Valine at this position leads to loss of function, which was captured by the MD/FEP simulations. The calculated ΔΔG_bind_ for alanine substitution was +0.918 kcal/mol and the experimental value was +0.85 kcal/mol. Structural insights in the mutated (alanine) state reveal the loss of water mediated interactions between the neighboring B9Ser and 91Glu, 59Arg from the Fn-III-1 domain, which may be the drivers of the observed loss-of-function. In the case of B10Val, the calculated ΔΔG_bind_ was -2.7 kcal/mol whereas the experimental value was +0.26 kcal/mol (Table S4). The obtained ΔΔG_bind_ value by considering full length human insulin was not in agreement with the experimental value. Since we had observed the effect of solvent-mediated and long-range interactions in our previous calculations, we re-performed this calculation using DesB30 insulin in line with experiment. The calculated ΔΔG_bind_ was 0.013 kcal/mol which was in-line with the experimental value. This calculation highlighted the ability of the FEP method to capture long-range effects, such as the deletion of B30 residue on ΔΔG_bind_ for this particular mutation, despite this change not being significant for multiple other variants.

#### B10Gln & B10Phe

Substitution of B10His with more polar amino acid Glutamine (Q) led to a small gain of function. In the case of B10Gln mutant, the calculated ΔΔG_bind_ was -0.359 kcal/mol and the experimental value was -0.36 kcal/mol (Table S4). Structural insights revealed a largely compensatory pattern of interactions, which included solvent mediated interactions between B13Glu and 91Glu of Fn-III-1 domain that was not observed in the WT-ins state. Similarly, for B10Phe, FEP has predicted near equipotency (ΔΔG_bind_ =0.13 kcal/mol) which was in line with experimental values (0.06 kcal/mol). Overall, recapitulation of the effects of different mutations at key B10 position along with the examination of the factors influencing the observed changes from both an energetic and structural perspective, elucidates the intricate interaction networks that govern the engagement of Insulin with IGF-1R at this specific position within the peptide.

### Understanding the gain of function associated with long-acting insulin glargine

Insulin glargine, marketed under the brand name Lantus® among others, is a long-acting human insulin analog designed to closely resemble the natural physiological profile of endogenous insulin.^33,56,57,57,58^ It differs from wild-type human insulin (WT-ins) by having two additional Arginine residues at positions B31 and B32 of the C-terminus of the B-chain, as well as a substitution of Asparagine with Glycine at position 21 of the A-chain. ^23,33,59^ These modifications contribute to its extended duration of action. Several factors contribute to the prolonged activity of insulin glargine, including its low solubility at physiological pH, a shift in isoelectric point from 5.4 to 6.7, and a low dissociation rate.^23,59,60^ While these factors play a significant role, there are other additional factors that also contribute to its long-acting profile.^23^

Insulin glargine undergoes proteolytic degradation leading to fully soluble and active metabolites, IM (Intermediate metabolite), M1 and M2.^23,61,62^ These metabolites have different sequence compared to insulin glargine. IM does not contain Arginine at B32 position, metabolite M1 is formed by loss of B31Arg, B32Arg, and metabolite M2 is formed by the deletion of B30Thr (DesB30) in comparison to insulin glargine.^23,61^ These metabolites are present in the plasma and partly contribute to the long-lasting effect of analog.^62^ The metabolic potency of insulin glargine relative to human insulin was ∼86% indicating the decrease in receptor binding to insulin receptor (IR).^20,23^ On the contrary, insulin glargine and metabolite IM were more potent and M1, M2 are less potent compared to human insulin, IGF-1 at IGF-1R receptor. ^23^

We wished to use FEP to understand the molecular insights for the observed mitogenic potencies for glargine, IM and metabolites M1 & M2. The FEP was performed using SBC in an similar setup to that used to study Insulin B10 analogs described in the previous section. Studying insulin glargine using FEP is a huge task owing to the differences in the number of residues, between the initial state (insulin glargine) to final state (insulin). Since FEP is a state function, schemes were carefully designed in order to include all the intermediate metabolites of insulin glargine to the initial state (WT-ins) and FEP scheme is provided in Figure S2. FEP calculations were performed using the final snapshot obtained from the equilibration step of MD simulations starting from the insulin-bound EM structure of IGF-1R [PDB accession code: 6JK8].^10^ In total 2.1 μs simulations were performed for all the four states (glargine, IM, M1 & M2) with an average of ∼280 ns used for each FEP calculation at the receptor. To determine the relative binding free energies (ΔΔG_bind_) of each mutant, thermodynamic cycle was constructed, with free ligand in water serving as the reference state (Figure S1b). The resulting ΔΔG_bind_ values were then compared to that of the WT-ins to measure the relative affinities of each molecule at the IGF-1R. The ΔΔG_bind_ obtained from the FEP calculations was in good agreement with the experimentally observed mitogenic potencies in all the four cases ( glargine, IM, M1 & M2-100 %) (Table S4).

In the case of insulin glargine, the calculated ΔΔG_bind_ value (-6.3 kcal/mol) correctly predicted gain of function compared to WT-ins with a significant overestimation compared to experimental value -1.27 kcal/mol. Decomposition of ΔΔG_bind_ revealed that the effect of electrostatic changes was not compensated by the annihilation of the arginine atoms. Structural analysis further revealed the interactions of B31Arg with residue of 305Gln and 286Ser of CR domain (Figure 6b) in the glargine state which were absent in WT-ins caused a gain of function. A part from these direct interactions, there were multiple mediated interactions between these Arginines and residues from L1, CR domain which were not found in the WT-ins. For the metabolite IM, the calculated ΔΔG_bind_ value was -5.4 kcal/mol, which was overestimated compared to the experimental value of -0.7 kcal/mol.^23^ Structural insights reveal the direct and solvent mediated interactions of B31Arg, B30Thr with 289Glu of CR domain lead to the gain of function for this metabolite.

**Figure-5:**
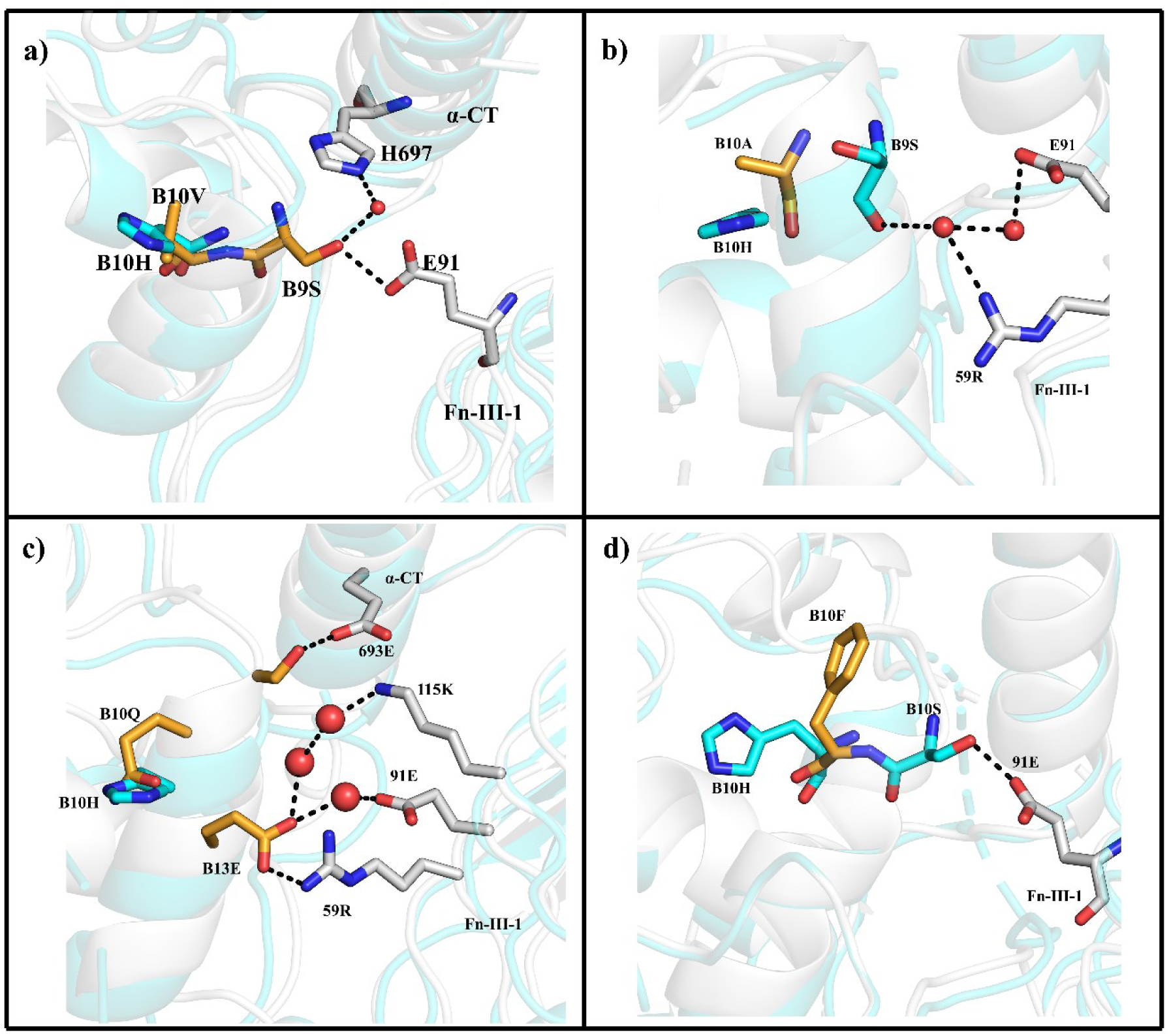
Structural analysis of B10Val, Ala, Gln, and Phe are shown compared to WT-ins. The WT-ins are represented by cyan color, mutant type in orange color. The interacting residues of receptor with insulin are colored in white color stick representation. Water oxygen atom is represented in red color. a) B10Val DesB30 system was shown, with interacting residues of B9Ser with 91Glu. b) In the B10Ala system, solvent mediated interactions between B9Ser and 54Arg, 91Glu were shown. c) In B10Gln system, the interactions between 59Arg, B13Glu, solvent mediated interactions B13Glu, 115Lys, 91Glu were shown, B13Glu interactions with 59Arg, 91Glu. d) B10Phe system, showing the interaction of B10Ser with 91Glu in WT-ins.

**Figure 6:**
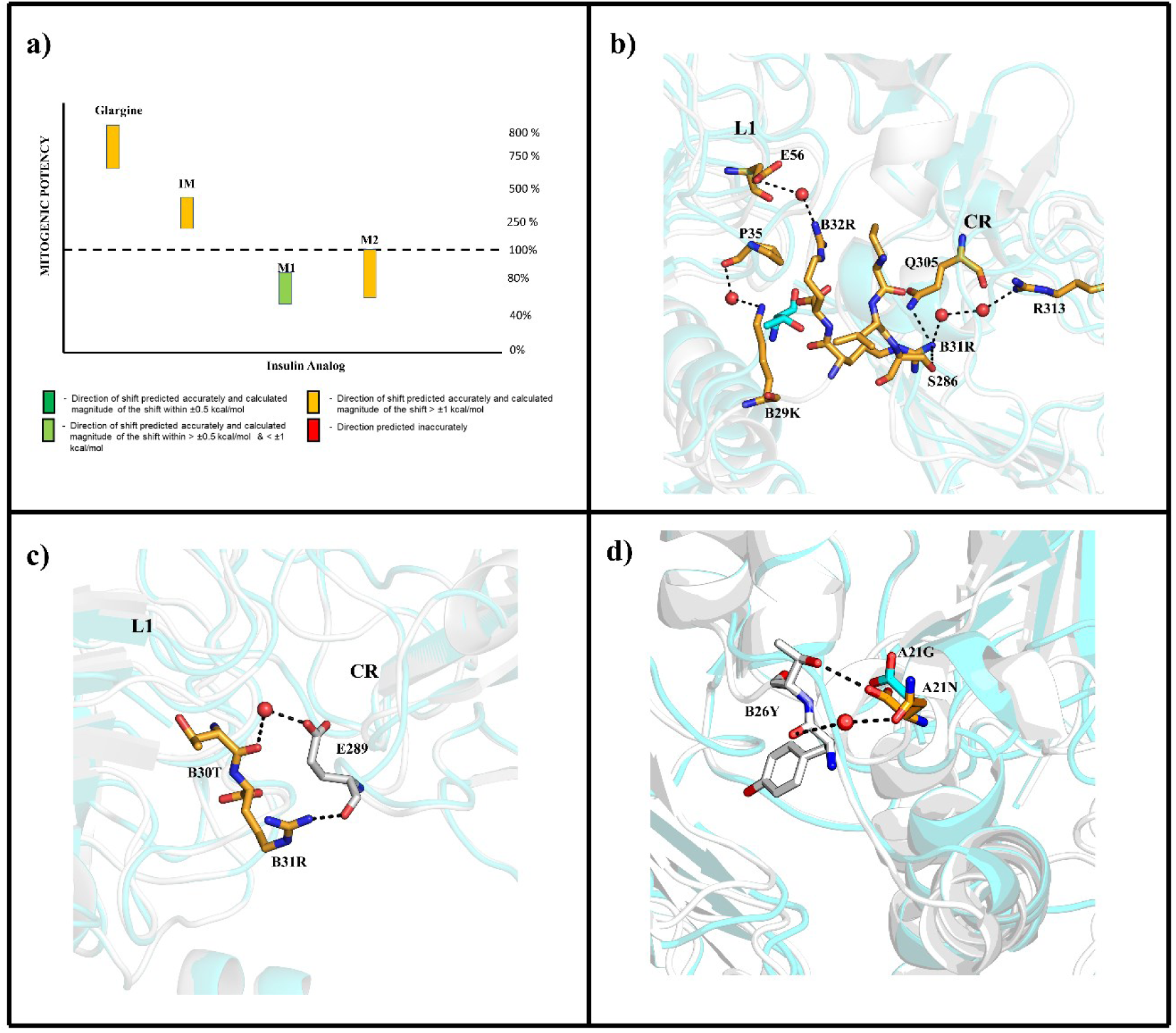
Structural analysis of insulin glargine and its metabolites. In all figure panels WT-ins complex is shown in cyan color and mutant in the orange color. Interacting residues are represented in sticks, insulin residues were colored in orange, receptor residues were shown in white color stick representation. a) correlation of mitogenic potency values from FEP and experiments, b) in glargine the interactions between B31Arg and 305Gln, 286Ser of CR-domain were shown, c) IM metabolite, interaction between B31Arg, B30Thr with 289Glu of CR domain were shown. d) In M1 metabolite interactions between A21Gln and B26Tyr were shown.

In the case of the metabolite M1 the calculated ΔΔG_bind_ value (0.46 kcal/mol) was in-line with the subtle loss of function observed at the IGF-1R (experimental value of 0.17 kcal/mol).^23^. The calculated ΔΔG_bind_ values for the metabolite M2 was +2.1 kcal/mol which overestimated the loss-of-function compared to experiment 0.23 kcal/mol. Decomposition of ΔΔG_bind_ revealed that the electrostatic interactions are more than compensated by the annihilation of Threonine atoms. Promisingly, despite the large perturbations attempted in these calculations, the correct rank-ordering of Insulin Glargine and its metabolites IM, M1 and M2 was recapitulated albeit with overestimations of potency shifts in a few cases.

## Discussion

IR and IGF-1R belong to the receptor tyrosine kinase family and share significant sequence similarity. Despite this similarity, the binding of their respective cognate ligands results in different functions. However, these ligands can also cross-bind to non-cognate receptors with lower affinity; for instance, insulin binds to IGF-1R approximately 100 times less effectively than IGF-1. The binding of insulin to IGF-1R has been associated with oncogenicity concerns, therefore, it is crucial to design new insulin analogs with low binding affinity at the off-target IGF-1R, to enhance the safety profile of these analogs. Although, discovery of insulin is more than a century old molecular level insights of its interactions with off-target IGF-1R remains elusive. In this study, we have leveraged the recent insulin/IGF-1-IGF-1R structures and used MD/FEP to elucidate ligand engagement with the IGF-1R using these complexes. Three key results emerged from our calculations. First, MD simulations captured the stability of bound insulin/IGF-1 at IGF-1R and revealed key long-lived interactions of cognate and non-cognate ligands with IGF-1R. This framework enabled us to assess and comprehend the recognition of insulin at its off-target receptor and understand the peculiarities associated with the inverted “V” and “T”-like bound conformations that have been observed upon ligand binding to the IGF-1R. Next FEP could capture the increase in potency of superactive insulin X10 and other analogs of insulin that varied at the B10 position. The molecular insights gleaned from these calculations could provide ways of leveraging changes in this position without causing significant increases in off-target binding. Finally, FEP accurately recapitulate the efficacy profiles for insulin glargine and its metabolites. The impressive ability of FEP to accurately rank-order glargine and its metabolites in terms of their mitogenic potencies highlighted the promise of this method to assess selectivity of insulin analogs.

In this study we have used recent structures of ligand-bound IGF-1R complexes to perform MD simulations.^2,10^ The structures to-date have revealed the presence of two distinct ligand bound conformations, namely an inverted “V”- or “T”-like shapes. The short MD simulations confirmed the overall stability of the cognate/non-cognate receptor complexes irrespective of the starting conformation. In the “T”-like insulin-bound conformation arising from the EM - structure, the RMSD is higher compared to the previous XRD structure which is attributed to the interaction N-termial A-chain residues and B14-B18 residues with L1 domain residues. Analysis of long-lived contacts considering 4.5 Å around the sphere indicate the importance of α-CT domain which is in line with the previous studies.^2,10^ It was observed that upon binding of IGF-1 to IGF-1R the 701Phe from receptor displaces 24Tyr, 25Phe residues of ligand, which further interacts with 702Val and 704Arg. This provided a rationale for how mutation of these residues leads to decrease in IGF-1 binding.^2,63,64^ For example, mutation of 704Arg residue of the IGF-1R receptor with Ala leads to decrease of IGF-1 binding by four times compared to wild-type receptor.^65^ The equivalent residue in insulin is 19Tyr of A-chain which also involved in similar interactions with the non-cognate receptor.^63,66^ Overall these simulations provided us with key interactions involved in the cognate/non-cognate complexes and could provide us with a framework to better understand the distinct engagement pattern of insulin and IGF-1 at the IGF-1R.

While the brief MD simulations offered a general overview of insulin/IGF-1 recognition at IGF-1R, comprehending the alterations in the functional profile resulting from the mutation/additions of specific residues is crucial for developing a deeper understanding of receptor-ligand interactions. As a first step, we wanted to understand substitutions at the B10 position of insulin which created lot of interest owing to the potential to increase metabolic potency via specific mutations. Substitution of B10His with aspartic acid gives “super-active” insulin which has significantly improved metabolic potency compared to human insulin (∼375 %).^19,25,27^ However, the even more significant potency jump at the off-target IGF-1R limited the usage of B10Asp owing to safety concerns.^20,22,27,54^ Understanding the molecular drivers behind the mitogenic potency profile of B10Asp could help us to improve specificity of analogs carrying an acidic amino acid at this position. The MD-FEP calculations could accurately capture the increase in potency associated with B10Asp. Interestingly, it was observed at this position that B10 was found to be completely solvent exposed at the IGF-1R with no direct interaction with the receptor. Upon examining the solvent networks surrounding the residue it was found that the shift from histidine to aspartic acid facilitated a strong network of water-mediated interactions, fostering inter-domain solvent bridges that stabilized the overall bound conformation at the IGF-1R. The solvent networks analysis highlighted the power of MD/FEP in not only capturing the energetics accompanying a mutation but further deconvoluting the contributions of different interactions towards the observed shifts in affinity. The simulations also showcased how the explicit inclusion of water molecules can capture significant effects that might not be immediately evident from static structures or even continuum solvation methods. The enhanced understanding gained from these calculations could prove invaluable in the design of novel analogs with tailored properties.

Since the position B10 in insulin is one of interest, we studied different amino acids varying in nature from hydrophobic (Ala/Val/Phe) to hydrophilic (Gln) at this position. FEP could capture the loss of function upon mutating this position with hydrophobic residues which is attributed to the loss of insulin mediated interactions between α-CT, Fn-III-1 domains. Substitution with the polar glutamine led to the forming of new interactions between different domains, which may be the driver behind the subtle gain in potency at IGF-1R. In the case of B10Val, an interesting result was obtained. The calculations with full length B10Val indicated a sharp increase in potency accompanying this mutation, which was contrary to experiment. Since the experimental mitogenic potencies were obtained using DesB30 systems, we repeated the calculation using DesB30 insulin, which resulted in values in-line with experiments. This result potentially showcases the ability of FEP method to capture long-range effects arising from changes distant from the immediate neighbourhood of the mutation. It would be interesting to see if full-length B10Val indeed demonstrates an increased mitogenic potency when compared to human insulin. Overall FEP could accurately capture the effect of various amino acids at the B10 position further showcasing its promise as a tool for rational design of selective insulin analogs.

Insulin Glargine is a long-acting insulin analogue marketed by Sanofi-Aventis under the brand name Lantus®.^56^ Insulin glargine disassociates into metabolites named as IM, M1, M2.^22^ Whereas Glargine and IM are associated with increased mitogenic potency, M1 and M2 have lower activities at IGF-1R when compared to human insulin. Studying glargine and its metabolites pushes the boundaries of perturbation theory and FEP due to the differences in sequence length compared to the reference human insulin. An elaborate FEP scheme was devised to capture the complexity associated with these calculations. Promisingly, FEP recapitulated the rank ordering of Insulin Glargine and its metabolites IM, M1 and M2 with respective to mitogenic potency. Furthermore, the interaction analysis provided a basis for understanding the significant increases in mitogenic potency for glargine and IM compared to human insulin. The overestimation in terms of magnitudes of shift potentially arose from insufficient reorganization accompanying the deletion of the large positively charged residues.^67^ The ability of FEP to capture the change in potencies in the face of such large changes highlighted the promise of this method as a generalized tool for the in silico evaluation of mutations including those involving change in sequence length.

## Conclusions

The current study delves into the molecular intricacies of cognate and non-cognate complexes at IGF-1R, thereby shedding light on the observed shifts in mitogenic potency within insulin X10, other insulin B10 mutants, insulin glargine and its metabolites. It illustrates the robustness of the FEP method in accurately capturing changes in free energy associated with a variety mutations at the off-target receptor, even identifying subtle alterations in long-range interactions within the structure. Notably, the study not only recaptured the functional profiles of different mutants but also disentangled free energy changes into individual contributions, enhancing the molecular level understanding of insulin engagement at IGF-1R. In essence, this study demonstrated the power of MD/FEP as a versatile tool for systematically evaluating mutations *in silico*, facilitating the rational design of selective novel insulins and peptides in general.

## Methods

### Homology modelling

Homology modeling is a computational structure prediction technique and widely used to predict the three-dimensional structure of the protein from the sequence. Often in the crystal structures obtained using various experimental techniques, some of the residues, domains may be missing. Homology model is often used to fill the missing resides and obtain a three dimensional structure. In this study, missing residues were found in both the structures which were modeled using Modeler 9.25 by considering the respective protein as a template.^68^ In the XRD structure of IGF-1R (pdb id: 5U8Q) the missing α-CT domain was modeled using insulin α-CT as template. Thus obtained complete structure was used as a starting structure for MD/FEP simulations.

### Molecular Dynamics (MD) Simulations

MD simulations were performed with the initial coordinates obtained from the crystal structure & EM structure with PDB accession codes: 5U8Q and 6JK8.^46,66^ After building the receptor-complex model without missing residues, appropriate protonation states were assigned. Most probable protonation state for aminoacids at pH=7 was set. Assigning of appropriate protonation states is very crucial as it will effect the output free energy values. The correct histidine tautomeric state was assigned using visual inspection. Thus, obtained model was solvated using TIP3P water model and counter ions were added to neutralize the complex.^69^ Thus obtained system was minimized and equilibrated using restraints on heavy atoms of protein in NPT ensemble for 10 ns using OPLSAA^70^ force field in GROMACS^71^ simulation engine for EM-structure. Similarly, for crystal structure AMBER14ffSB^72^ force field was used and the simulations were performed using AMBER simulation engine.^73^ The constraints were applied using SHAKE algorithm.^74^ Long range electrostatics were treated using EWALD^75^ summation method. The Langevin thermostat was used to maintain temperature and Berendsen barostat^76,77^ was used to maintain pressure of 1 Bar with coupling constant of 2 ps. After equilibration, the system was subjected to 50 ns unrestrained simulation. The final snapshot from the equilibration MD simulation was used as input to compute free energy (ΔΔG) using FEP method using spherical boundary conditions (SBC).

### MD using SBC and the Free Energy Perturbation (FEP) Method

FEP method was used to compute free-energy differences for an alchemical transformation of from an initial state A to final state B.^78–80^ The free energy (ΔΔG_bind_) were computed using FEP implemented using Q program.^81^ In our study, the initial state refers to the mutant state and the final state refers to the insulin state. The input structure for the FEP simulations was obtained from the last snapshot from the equilibration MD simulations. Unlike MD simulations, the FEP calculations were performed using spherical boundary conditions (SBC). A representative image of PBC to SBC conversion was shown in the Figure S1a. In the current study, we have chosen Cδ atom of B15Leu of insulin as center and considered a sphere with radius 30 Å at IGF-1R. Heavy restraints were applied for the atoms outside the sphere in order to maintain the structure close to the initial structure.

All the protein, ligand and water atoms with the defined radius are explicitly considered. OPLSAA force field were used for the receptor complex. Thus obtained system was equilibrated in a series of six steps with varying temperature and restraints on the heavy atoms. Finally, the well equilibrated system was used as input to run unrestrained production run which was used to compute free energies. Atoms close to sphere edge were restrained to their initial coordinates and atoms beyond sphere edge were excluded from nonbonded interactions. Asp, Glu, Lys and Arg residues within radius of 30 Å are protonated according to their most probable state at pH 7 and ionizable residues close to sphere were neutralized. Residues lying outside the sphere are restrained to stay close to the observed structure, so that we can evaluate energy of this conformational region rather than allowing system to move into other conformational regions.^82^ The protonation state for Histidines within sphere were assigned based on visual inspection. The SHAKE algorithm was used to constraint all the bonds, all angles and water molecules at the sphere surface were subjected to radial and polarization restraints according to the SCAAS model.^74,77,81^ Long range electrostatics were treated with local reaction field (LRF) method.^83^ The time step was set to 1 fs and non-bonded pair lists were updated every 25 steps.

The following FEP protocol was carried out to obtain relative binding free energy for the alchemical transformations of the residues: i) the transformation of partial charges and ii) combined transformation of Lennard-Jones (LJ) and parameters involving covalent bonds in several MD/FEP calculations. To annihilate heavy atoms, a separate MD/FEP calculations were performed to remove them in step-wise manner. A softcore potential was introduced for the atom in a first step and followed by removal of the resulting van der Waal’s potential. The force field parameters describing angles, bonds, and proper/improper dihedrals were retained for annihilated atoms. A summary of the FEP schemes was provided in Figure S2. Each FEP calculation were divided into *n* intermediate states which were equilibrated parallelly. The potential (U_m_) for the respective transformation A (initial state) and B (final state) is given by

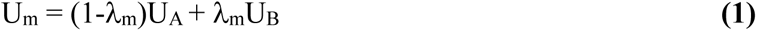

where λ_m_ varies from 0 to 1. A value of λ_m_=0 represents initial state, λ_m_=1 represents final state and values in between 0 and 1 represents intermediate states. The transformations involving partial charges were performed using up to 36 λ_m_ steps and the number of λ_m_ steps used to transform LJ and bonded parameters ranged from 42 to 63 λ_m_ steps depending upon the complexity of the transformation. The ligand-receptor complex was typically equilibrated for 1.76 ns at each λ_m_ step. During the equilibration of the system the harmonic restraints were released on ligand-receptor complexes in several steps and the temperature was increased in a series of steps to 310 K. After equilibration, the unrestrained simulation for the complex was typically carried out for 0.5 ns for each λ_m_ step from which potential energies were extracted. The ligand in water consisted of the entire insulin molecule and was typically equilibrated for 0.76 ns followed by 0.25 ns of unrestrained simulation for each λ_m_ step. The same sphere radii were maintained between the respective receptor and water simulations. The difference in free energy between initial state (A) and final state (B) was calculated by summing up the free energy differences of the n intermediate states using the relation

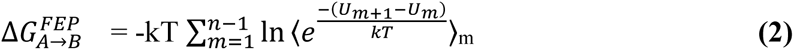

Where 〈… 〉_*m*_ represents an ensemble average on the potential U_m_ which is calculated from the MD simulations, k is Boltzmann constant and T is absolute temperature.^84^ The uncertainty of a transformation was quantified as the difference in free energy obtained by applying FEP formula in the forward and reverse direction and was minimized by increasing the number of λ_m_ values or simulation length until convergence was obtained.

### Water(solvent)-interaction network analysis

Solvent network plays a crucial role in stabilization of complexes. To examine the stabilization of initial state or final state through solvent mediated interactions, we have performed solvent networks analysis around the WT-ins and mutant. In this analysis, we have defined an interaction if there is an residue or water molecule present within 3.5 Å cutoff of reference residue. First order contacts are those interactions with the receptor that were within 3.5 Å of the residue of interest which was named as (P0), and if there is any water molecule within the same cutoff, it will be labelled as W0. Using the W0 waters as the new centers, the analysis was repeated. A receptor polar contact within 3.5 Å of any W0 water was defined as a second order contact (P1) and any water molecule within the same radius as a second-order water contact (W1), the process was repeated with W1 waters as centers resulting in a list of third-order receptor (P2) or water (W2) contacts. Average values of W0, P0, W1, P1, W2 and P2 for any residue of interest were obtained by averaging across the number of considered snapshots.

