## Supplementary file for "Mapping Structural Drivers of Insulin and its Analogs at the IGF-1 Receptor Using Molecular Dynamics and Free Energy Calculations"

Table S1**:** A summary of currently available XRD and Cryo-EM structures for IGF-1R receptors in complex with different ligands or as apo receptors.

| Receptor | | Method | PDB IDs | #SS | Year | Binding Site | Ref. |
|---|---|---|---|---|---|---|---|
| **IR/IGF-1R Hybrid** | IGF-1 | XRD | 4XSS | 3 | 2015 | - | ([1]) |
| **IGF-1R** | IGF-1 | XRD | 5U8Q<br>5U8R | 3 | 2018<br>2018 | - | ([2])<br>([2]) |
| **IGF-1R** | IGF-1 | EM | 6PYH | 3 | 2019 | HA | ([3]) |
| **IGF-1R** | Insulin (wild type) | EM | 6JK8 | 3 | 2020 | - | ([4]) |
| **IGF-1R** | IGF-1 | EM | 7YRR | 3 | 2023* | - | |
| **IGF-1R** | Insulin | EM | 7V3P | 3 | 2022* | | |

Note**:** #SS: Number of disulfide bridges in Ligand. BS: binding site type. HA: High affinity and LA: Low affinity

*: Indicates the structure deposited in the RCSB but the article is not published

Table S2**:** Binding enthalpies calculated from MM-GBSA calculation. Mean, SEM and Sd. are calculated from the ΔH calculated from ligand molecule bound at two HA sites, s1 & s1' from two simulations with different initial velocities and solvent accessible surface area (SASA) values for corresponding ligands.

| Complex | $\Delta H_{bind}$ (kcal/mol) (MM-GBSA) | | SASA($Å^2$) (LCPO, surface) | |
|---|---|---|---|---|
| | Mean | SEM | Mean | Std. Dev. |
| 5U8Q: IGF-1 | -71.46 | 1.67 | 678.39 | 65.50 |
| 5U8Q: WT-ins | -55.06 | 7.31 | 643.88 | 60.55 |

Table S3: Ensemble averaged solvent interaction networks for B10Asp & B10His in IGF-1R from MD trajectories.

| | Protein | | Water | |
|---|---|---|---|---|
| | IGF-1R | | IGF-1R | |
| | B10Asp | B10His | B10Asp | B10His |
| Zero order contacts | 0 (P0) | 0 (P0) | 6.68 (W0) | 2.74 (W0) |
| First order contacts | 0.24 (P1) | 0.08 (P1) | 25.94 (W1) | 9.98 (W1) |
| Second order contacts | 2.02 (P2) | 0.64 (P2) | 38.84 (W2) | 22.38 (W2) |

Table S4: Calculated relative free energies ($\Delta\Delta G_{bind}$) at IGF-1R calculated for all the X10, B10, insulin glargine and its metabolites considered in this study along with the range of experimental values available for each analog. The experimental spread of $\Delta\Delta G_{bind}$ has been calculated using the inner means and a mean of means (if more values are available) has been considered in order to compare with the calculated $\Delta\Delta G_{bind}$. The experimental and calculated $\Delta\Delta G_{bind}$ have both been reported in relation to WT-ins.

| Transformation | State-1 | State-2 | Experimental ΔΔG (mean)(kcal/mol) | FEP ΔΔG (kcal/mol) | Trend | Error (kcal/mol) (exp-sim) |
|---|---|---|---|---|---|---|
| B10Asp✓■ | Histidine | Aspartic acid | -1.26 | -2.98 | Yes (orange) | 1.7 |
| B10Ala✓ | Histidine | Alanine | 0.85 | 0.918 | Yes (green) | -0.03 |
| B10Val#✓ | Histidine | Valine | 0.26 | 0.013 | Yes (green) | 0.28 |
| B10Gln✓ | Histidine | Glutamine | -0.36 | -0.359 | Yes (orange) | -0.005 |
| B10Phe✓ | Histidine | Phenyl alanine | 0.06 | 0.13 | Yes (dark green) | -0.07 |
| Insulin glargine✓■ | TRR | Insulin (T) | -1.27 | -6.3 | Yes (orange) | 5.02 |
| IM✓■ | TR | Insulin (T) | -0.7 | -5.4 | Yes (orange) | 4.6 |
| M1✓ | N | G | 0.17 | 0.46 | Yes (green) | -0.2 |
| M2✓■ | N | COO- | 0.23 | 2.1 | Yes (orange) | -1.8 |

✓ Indicates the direction of shift predicted from FEP is in-line with experiments.

× Indicates the direction of shift predicted from FEP is not in-line with experiments

○ Indicates the magnitude of shift predicted from FEP is under estimated from the mean with a difference in magnitude of >1 kcal/mol

■ Indicates the magnitude of shift predicted from FEP is over estimated from the mean with a difference in magnitude of > 1 kcal/mol

#: Fep calculations were performed using DesB30 system.

The color below the deviation from experimental mean values (column trend) indicate the accuracy of the prediction. Dark green indicate that the direction of shift was predicted correctly and magnitude of difference between FEP and mean experimental values are < 1 kcal/mol. Orange indicates that the direction of shift predicted accurately and magnitude of difference between FEP and mean experimental values are > 1 kcal/mol. Red indicates the direction of shift predicted inaccurately by FEP. The same color scheme is followed in Figure 5 & 6 of the main text.

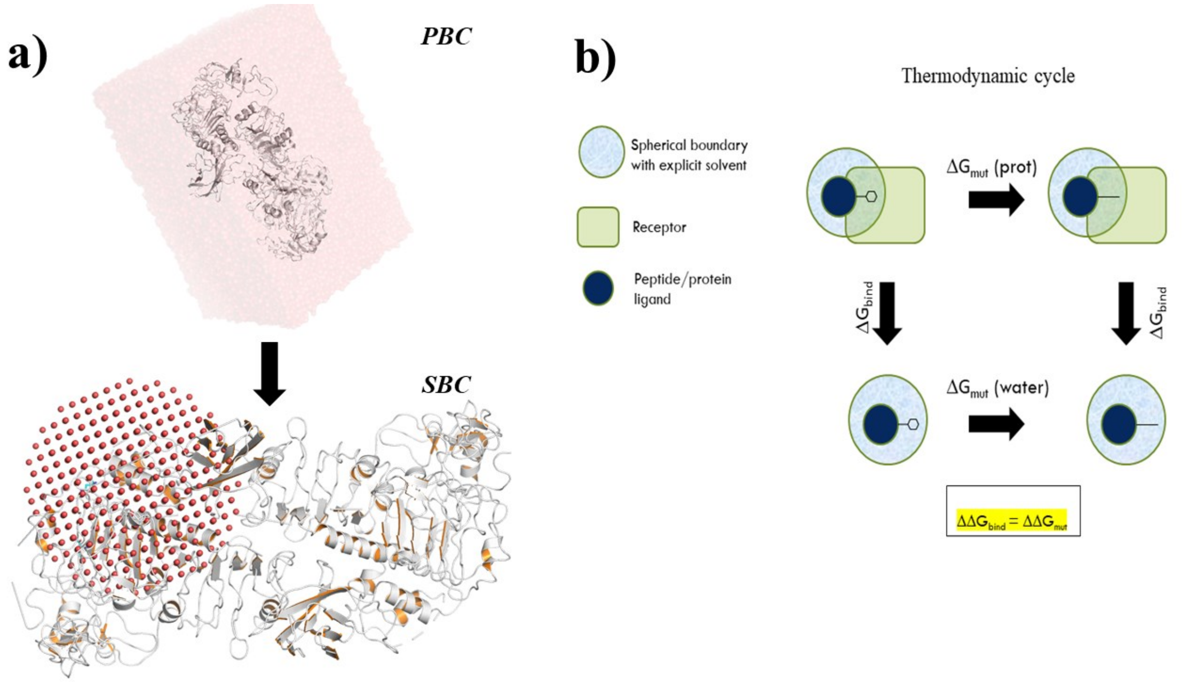


**Figure S1:** a) Depicts periodic boundary conditions (PBC) used for MD simulations and spherical boundary conditions (SBC) used for MD-FEP calculations. The top panel shows water molecules around the protein (cartoon) as surface representation in the PBC simulations. The bottom panel shows SBC used for FEP calculations in this study. All atoms within the sphere are fully flexible whereas those outside the sphere are restrained to their initial coordinates. The water molecules are shown as red spheres with receptors shown as cartoons.

b) Schematic representation of thermodynamic cycle used for FEP calculations. The blue circle represents the sphere, green rounded rectangle represents receptor and filled dark blue circle represents the peptide ligand. The same mutations are performed with the peptide bound to the receptor and with free peptide in water. The relative $\Delta\Delta G_{bind}$ is calculated by taking advantage of the thermodynamic cycle as shown.

**FEP scheme for insulin glargine**

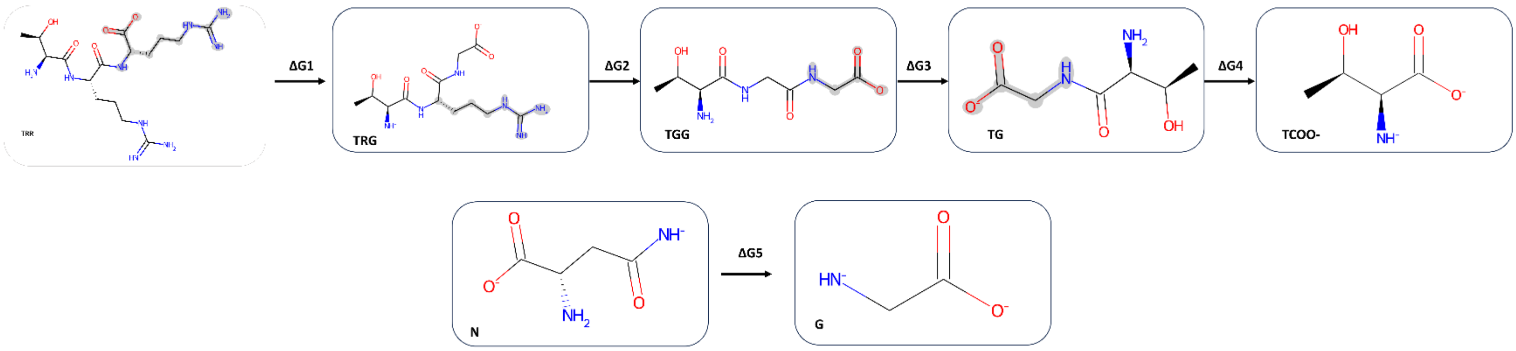


**Figure S2:** The above scheme was followed to carry out FEP transformation from insulin glargine as the starting state and insulin (final state). The below equations were used to compute ΔΔG.

$$\Delta\Delta G_{Glargine \longrightarrow WT\text{-}ins} = \Delta G1 + \Delta G2 + \Delta G3 + \Delta G4 + \Delta G5$$

$$\Delta\Delta G_{IM \longrightarrow WT\text{-}ins} = \Delta G2 + \Delta G3 + \Delta G4 + \Delta G5$$

$$\Delta\Delta G_{M1 \longrightarrow WT\text{-}ins} = \Delta G4$$
